# Genomic infectious disease epidemiology in partially sampled and ongoing outbreaks

**DOI:** 10.1101/065334

**Authors:** Xavier Didelot, Christophe Fraser, Jennifer Gardy, Caroline Colijn

## Abstract

Genomic data is increasingly being used to understand infectious disease epidemiology. Isolates from a given outbreak are sequenced, and the patterns of shared variation are used to infer which isolates within the outbreak are most closely related to each other. Unfortunately, the phylogenetic trees typically used to represent this variation are not directly informative about who infected whom - a phylogenetic tree is not a transmission tree. However, a transmission tree can be inferred from a phylogeny while accounting for within-host genetic diversity by colouring the branches of a phylogeny according to which host those branches were in. Here we extend this approach and show that it can be applied to partially sampled and ongoing outbreaks. This requires computing the correct probability of an observed transmission tree and we herein demonstrate how to do this for a large class of epidemiological models. We also demonstrate how the branch colouring approach can incorporate a variable number of unique colours to represent unsampled intermediates in transmission chains. The resulting algorithm is a reversible jump Monte-Carlo Markov Chain, which we apply to both simulated data and real data from an outbreak of tuberculosis. By accounting for unsampled cases and an outbreak which may not have reached its end, our method is uniquely suited to use in a public health environment during real-time outbreak investigations. We implemented our technique in an R package called TransPhylo, which is freely available from https://github.com/xavierdidelot/TransPhylo.

## Introduction

Infectious disease epidemiology is increasingly incorporating genomic data into routine public health practice, using genome sequencing for diagnosis, resistance typing, surveillance, and outbreak reconstruction. In the latter use case, we can draw inferences about the order and direction of transmission based on the presence of mutations common to multiple pathogen isolates (Gilchrist et al., 2015; Croucher and Didelot, 2015). While early works in this area assumed that pathogen genomes from a transmission pair should be identical or near-identical, a number of genomic outbreak investigations revealed the complicating factor of within-host evolution (Ypma et al., 2013; Romero-Severson et al., 2014; Worby et al., 2014).

Many important bacterial pathogens have periods of latency, chronic infection, or prolonged asymptomatic carriage, all of which contribute to the generation of within-host genetic diversity (Didelot et al., 2016). *Staphylococcus aureus* is a canonical example, in which single hosts can harbour multiple distinct lineages of the pathogen, each of which may be transmitted onwards (Young et al., 2012; Golubchik et al., 2013; Harris et al., 2013; Tong et al., 2015; Paterson et al., 2015; Azarian et al., 2016). In scenarios where a single host harbours substantial diversity, it can be difficult to infer which other hosts they infected - different lineages may have been transmitted at different points during the donor’s infection and the genome sequenced from the donor may only represent a single lineage captured at the time a diagnostic sample was taken and not the complete set of lineages present within that individual. Indeed, simulation studies have shown that if within-host diversity is ignored, incorrect inferences can be drawn about the transmission events that occurred within an outbreak (Romero-Severson et al., 2014; Worby et al., 2014; Worby and Read, 2015).

We have previously introduced a framework for inferring person-to-person transmission events from genomic data that considers within-host genetic diversity (Didelot et al., 2014). We use the genomic data to build a time-labelled phylogeny, which we divided into subtrees, each of which captures the variety of lineages that were present within each host. In other words, the phylogeny is “coloured” with a unique colour for each host, with transmission events represented as changes in colours along a branch. We originally used a simple susceptible-infected-recovered (SIR) model to evaluate the probability of the transmission tree, and we recently showed we can extend our approach to incorporate other types of epidemiological models (Hatherell et al., 2016). A similar approach, developed independently (Hall et al., 2015), couples phylogeny construction and transmission tree inference into a single step.

The main limitation of these previous methods is that they assume that all outbreak cases have been sampled and sequenced and that the outbreak has reached its end. These assumptions greatly simplify transmission tree inference, but don’t reflect epidemiological reality. An outbreak is rarely completely sampled - cases may not be reported to public health or they may not have nucleic acid available for sequencing - and genomic epidemiology investigations are frequently unfolding in real-time, meaning an outbreak is being analysed before it is ended. The few methods that can deal with unsampled cases do so at the cost of assuming no within-host diversity (Jombart et al., 2014; Mollentze et al., 2014). Here, we introduce a new Bayesian method for inferring transmission events from a timed phylogeny that can be applied to outbreaks that are partially sampled, ongoing, or both. We solve two problems that arise from these sampling issues: the complexity of calculating the probability of an observed transmission tree under these conditions, and the difficulty in exploring the posterior distribution of possible transmission trees given a phylogeny. Our method also permits the inference of when these transmission events occurred; when coupled with the person-to-person inference, this results in a comprehensive and epidemiologically actionable outbreak reconstruction. Here, we apply our new approach to both simulated datasets and a real-world dataset from the genomic investigation of a tuberculosis outbreak in Hamburg, Germany.

## Methods

We use a two-stage approach, first constructing a timed phylogenetic tree 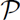 on the observed sequences and overlaying transmission events (Didelot et al., 2014). Let 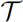 be the transmission tree, 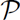 be the timed phylogenetic tree, *θ* be the parameters of the transmission and sampling model, and *N_e_g* the within-host effective population size.

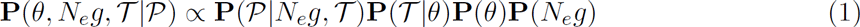

We compute 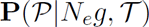 by separating 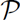 into independent parts, each of which evolves in a different individual host (Didelot et al., 2014; Hall et al., 2015); see below. This separability relies on the assumption of a complete transmission bottleneck, meaning that that within-host genetic diversity is lost at transmission, as is commonly assumed in this context. The central challenge here is therefore to compute 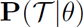 for a general model of transmission: one that allows for both unsampled cases and varying levels of infectivity throughout the course of infection, which is representative of the biological reality for many pathogens. We first illustrate how to do this in a scenario where the outbreak is over; this is a convenient assumption mathematically and makes the derivation simpler. We then proceed to the case where data collection ends at a fixed time before the end of the outbreak, as is the case when analysing an ongoing outbreak.

### Basic epidemiological model

The epidemiological process we consider is a stochastic branching process in which each infected individual transmits to secondary cases called offspring (Becker, 1977; Farrington et al., 2003). The number of offspring for any infected individual is drawn from the offspring distribution *α(k)* and we follow previous studies (Lloyd-Smith et al., 2005; Grassly and Fraser, 2008) in assuming that it is a negative binomial distribution with parameters (*r, p*). The mean of this distribution is called the reproduction number (Anderson and May, 1992), which is constant and equal to *R = rp/(1 — p)*, and the probability of having *k* offspring is 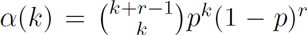. The time span between the primary and any secondary infection is drawn from the generation time distribution γ(τ), where τ is the time since infection of the primary case. The generation time distribution can take any form (Fine, 2003) but a Gamma distribution is often used (Wallinga and Lipsitch, 2007).

### Finished outbreak scenario

We first consider the situation where an outbreak follows the model above until there are no more infected individuals; we refer to this as a finished outbreak and we use the star subscript (*) to denote the mathematical quantities associated with this scenario. In this situation, all individuals are sampled with the same probability π, in which case the time span from infection until sampling has distribution *σ(τ)*. We want to calculate the probability of a transmission tree 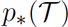. This requires some preliminary quantities.

We say that an infected individual is “included” if they are part of the transmission tree by being either sampled or by leading through transmission to at least one sampled descendant. Otherwise, we say that an infected individual is “excluded”. Let *ω_*_* be the probability of being excluded. This means the individual and all of their descendants are unsampled. Considering the number of offspring *k*, we have that:

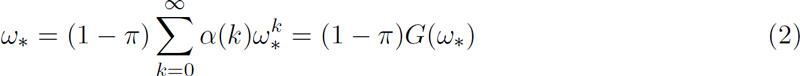

where *G(z)* is the probability generating function of the offspring distribution. We model this as a negative binomial distribution so that 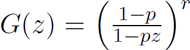, but our approach could use other distributions. We choose the negative binomial distribution to allow individuals to have different rates at which they are in contact with others (gamma-distributed) combined with a Poisson distribution of secondary infections given their individual rate. The solution *ω_*_* to Equation 2 is calculated numerically (Supplementary Material).

The probability that an individual has *d* offspring who are included in the process is

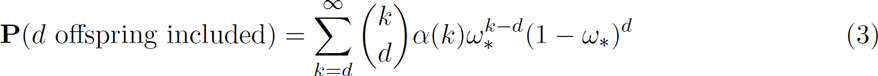

In our final product for 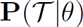, arrived at by induction, each included case will have its own term. For notational simplicity we define the “modified offspring function” to collect the other parts of this expression:

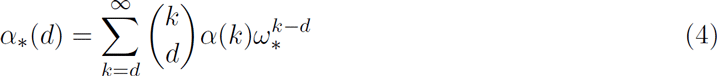

A good approximation is obtained by taking the sum up to a large value (Supplementary Material). Note that if *π* = 1 then *ω_*_* = 0 and *α_*_(d)* = *α(d)*.

We now consider a transmission tree 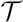 generated from the model, which is made of *n* nodes corresponding to the included infected individuals (either sampled or unsampled). They are indexed by *i* = 1,…,*n*. Let *s_i_* = 0 if *i* is unsampled and *s_i_* = 1 if *i* is sampled, in which case its sampling time is 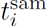. Let 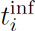 denote the time when *i* became infected and *d_i_* denote its number of included offspring who are indexed by j = 1..*d_i_*. The probability of 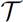 given the parameters *θ* can be obtained by considering the root *ρ* of the tree which has *d_ρ_* offspring, and the subtrees 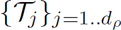 corresponding to each offspring. A recursive form of the probability of the transmission tree can then be written as:

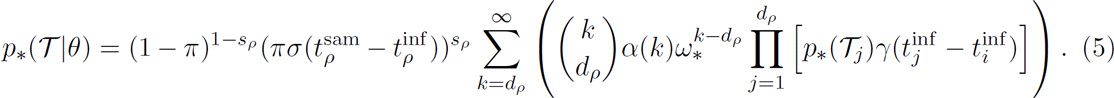

The parameters *θ* appear in the offspring distribution *α*, the generation time density γ and the sampling time density *σ*. The terms in the square brackets do not depend on *k*, so that we can rearrange the equation using the modified offspring function *α_*_* defined in Equation 4:

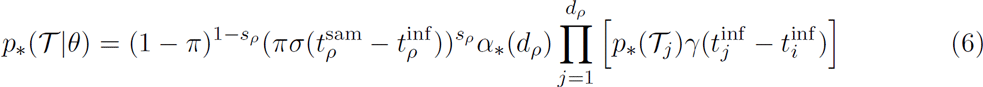

Finally by induction we obtain the probability of 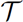 as a product over all nodes of the transmission tree:

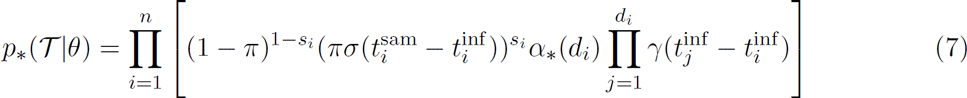

### Ongoing outbreak scenario

We now consider the situation where an outbreak follows the same model as previously described, until some known time *T* where observation stops. Whereas individuals were previously all sampled with the same probability *π*, it is now necessary to account for the fact that individuals who became infected soon before *T* have a lower probability of being sampled. More formally, the probability of sampling for an individual infected at time *t* is equal to:

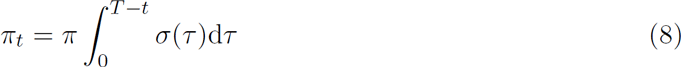

Stopping observation at time *T* also affects the probability of being excluded, with all individuals infected at *t ≥ T* being excluded.

For an individual infected at time *t*, let *ω_t_* be the probability of being excluded. Note that where *t > T*, *ω_t_* = 1. Before that time, *ω_t_* is not constant, but we know that as *t → −∞*, we should have *ω_t_ → ω_*_*. We have that:

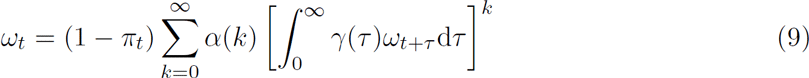

Let 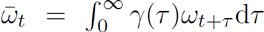. Using the generating function *G(z)* of the negative binomial distribution of *α(k)* we have 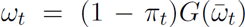. We approximate 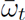 using a numerical integration (Supplementary Material). Good agreement is found with the expected limit *ω_−∞_ = ω_*_* where *ω_*_* is given in Equation 2.

As before, we use the modified offspring function to simplify the notation:

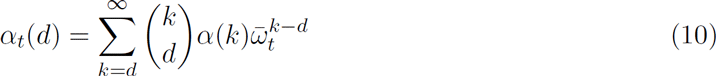

and obtain a good approximation by taking the sum up to a large value of *k* (Supplementary Material).

With the same recursive reasoning as in the finished outbreak scenario, we have:

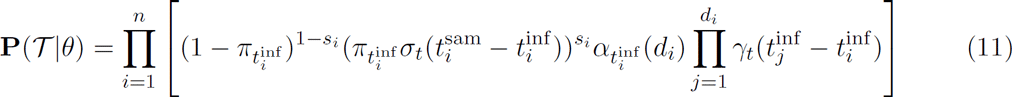

where *σ_t_(τ)* and *γ_t_(τ)* are respectively equal to *σ(τ)* and *γ(τ)* truncated at time *τ = T - t*.

### Inference of transmission tree given a phylogeny

The models described above generate transmission trees where each node is an infected individual, each terminal node is a sampled infected individual, and links between nodes represent direct transmission events (Figure 1A). Let us now consider that transmission involves the transfer of only a single genomic variant of the pathogen from the donor to recipient (ie a complete transmission bottleneck) and that sampling involves sequencing a single genome, randomly selected from the within-host pathogen population. The ancestry of the sequenced genomes can then be described as a phylogeny which is made of several subtrees, each of which corresponds to the evolution within one of the included hosts and describes the ancestral relationship between the genomes transmitted and/or sampled from that host (Figure 1B). The probability 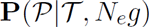 of a pathogen phylogeny 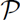 given a transmission tree 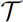 and within-host effective population size *N_e_g* is therefore equal to the product of the subtree likelihoods for all included hosts (Didelot et al., 2014; Hall et al., 2015), which can be calculated for example under the coalescent model with constant population size *N_e_g* (Kingman, 1982; Drummond et al., 2002).

**Figure 1.**
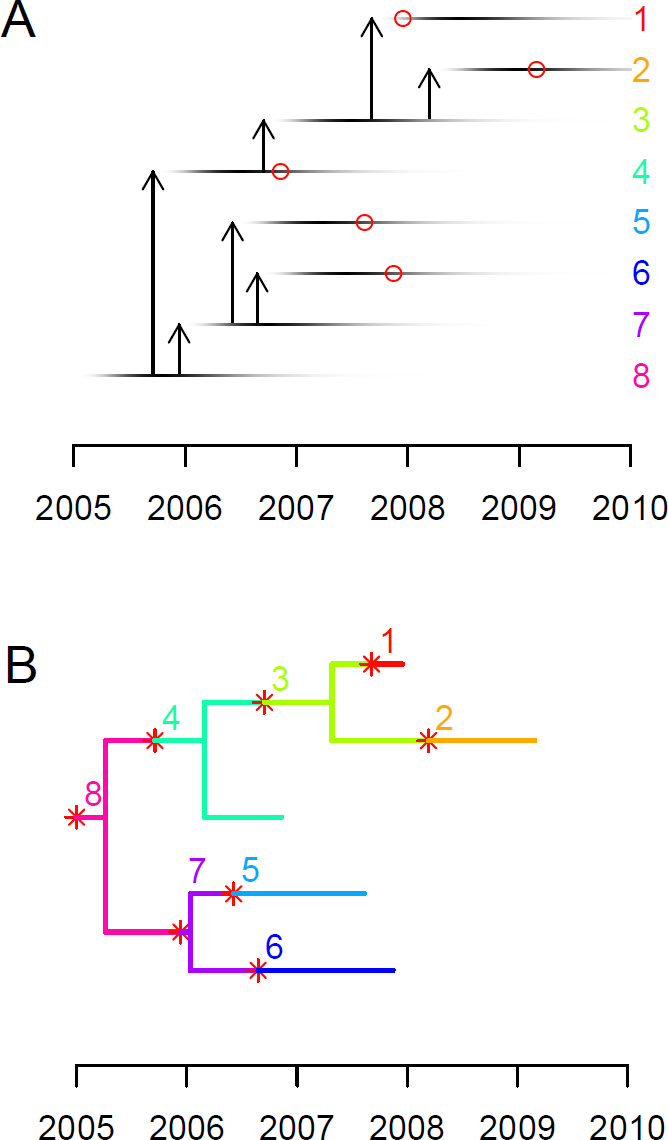
A: An illustrative example of transmission tree, with each horizontal line representing a case, and the darkness of each point representing their changing infectivity over time. Vertical arrows represent transmission from case to case. The red circles indicate which individuals were sampled (1, 2, 4, 5 and 6) and when. B: An example of colored phylogeny which corresponds to the transmission scenario shown in part A. Evolution within each host is shown in a unique color for each individual, as indicated by the labels and on the righthand side in part A. Red stars represent transmission events and correspond to the arrows shown in part A. Tips of the phylogeny represent sampled cases as shown by the red circles in part A.

Having defined both 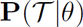 and 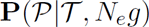, we can now perform Bayesian inference of the transmission tree 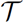 given a timed phylogeny 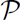 using the decomposition in Equation 1. Although a timed phylogeny is not directly available, there are powerful approaches readily available to reconstruct it from genomic data (Drummond et al., 2012; Bouckaert et al., 2014; Biek et al., 2015; To et al., 2016). As in our earlier work (Didelot et al., 2014), we can approach this problem by coloring the phylogeny with one color for each host (Figure 1B); however, since we now consider that some hosts may not have been sampled, the number of infected hosts and therefore the number of colors is not known. In other words, the parameter space is not of fixed dimensionality, and exploring it with a Monte-Carlo Markov Chain (MCMC) requires that we include reversible jumps that change the number of hosts in the transmission tree (Green, 1995). Our proposal for adding new transmission events is uniformly distributed on the edges of the phylogeny 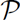. Our proposal for removing transmission events is uniformly distributed on the set of transmission events that can be removed without invalidating the transmission tree. In a transmission tree 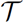 with *n* hosts and 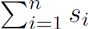 sampled hosts, there are 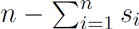 such removable transmission events. The Metropolis-Hastings-Green ratio for the MCMC move from 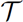 to 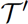 by adding a transmission event is therefore equal to:

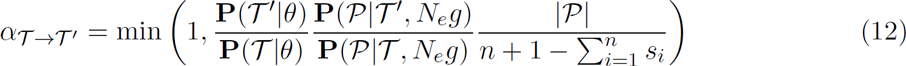

where 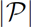 denotes the sum of the branch lengths of the phylogeny 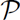. Conversely, the acceptance ratio of the MCMC update from 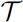 to 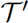 by removing a transmission event is:

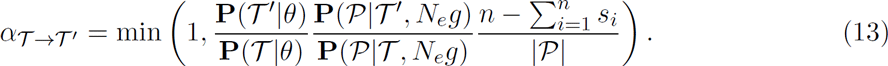

Within each MCMC iteration, additional standard Metropolis-Hastings moves are used to estimate the first parameter *r* of the Negative binomial distribution for the number of offspring (using an Exponential(1) prior), the second parameter *p* of the Negative binomial distribution of the number of offspring (using a Uniform([0,1]) prior), the probability of sampling π (using a Uniform([0,1]) prior), and the within-host effective population size *N_e_g* (using an Exponential(1) prior).

## Results

### Example application to a simulated dataset

We simulated an outbreak in which the generation time distribution had a gamma distribution with a mean of 1 year, with a negative binomial offspring distribution with parameters (*r* = 2, *p* = 0.5), such that the reproduction number was *R* = 2. We set the sampling density at π = 0.5 with a sampling time distribution identical to the generation time distribution. The simulation was stopped after *n* = 100 genomes had been sampled, which happened at time *T*. The corresponding phylogeny (Figure 2A) was used as input for our transmission tree inference algorithm with the date *T* used as described in the “ongoing outbreak scenario” in the Methods section. Performing 50,000 MCMC iterations took less than an hour on a standard computer. The mean posterior of the sampling proportion π was 0.48 with a 95% credibility interval of [0.36,0.59]. The mean posterior of the reproduction number R was 2.168367 with a 95% credibility interval of [1.75,2.65]. The estimates of these two key parameters of our model are therefore in excellent agreement with the true values used to perform the simulation.

**Figure 2.**
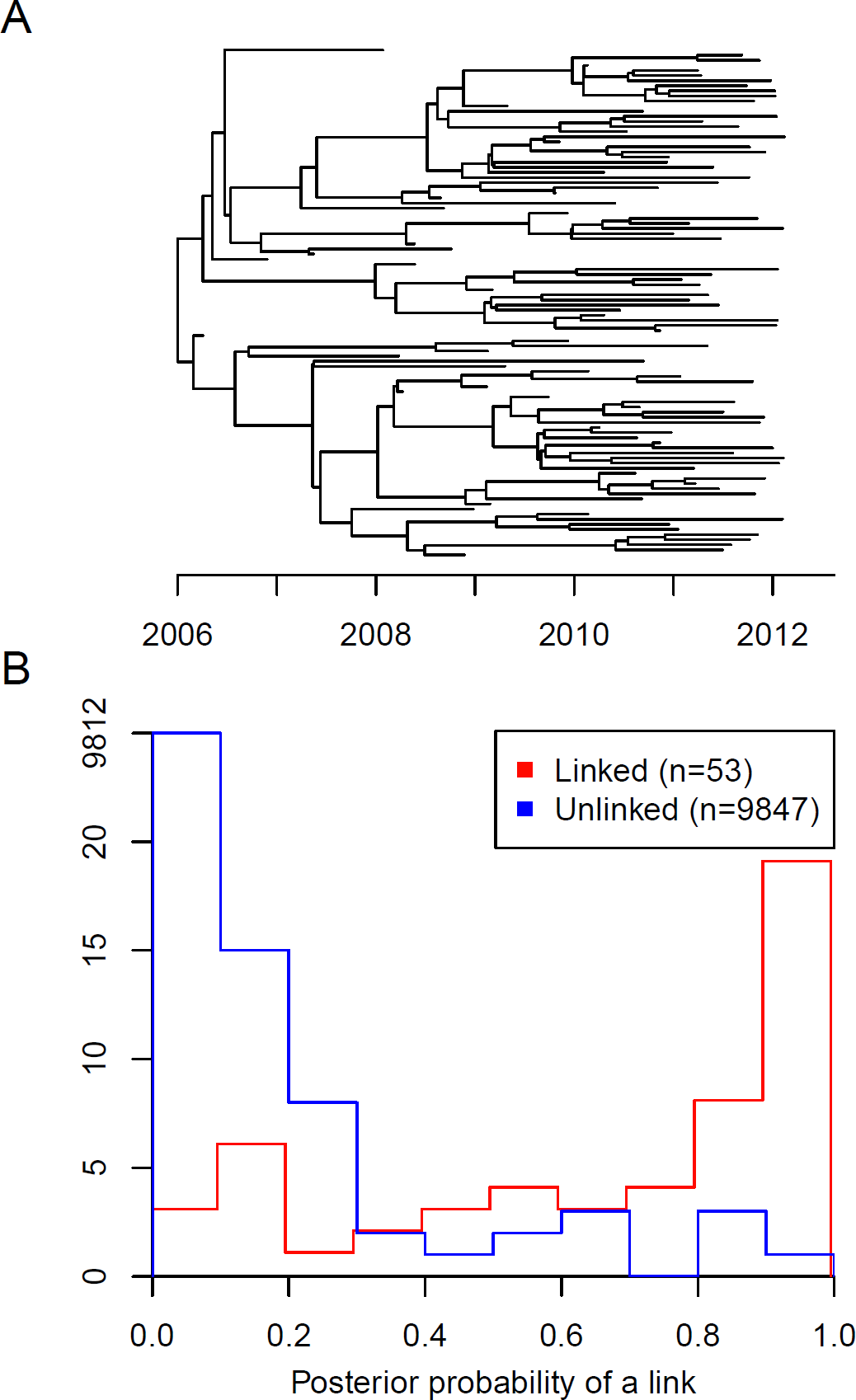
A: Timed phylogeny showing the relationship between 100 genomes sampled with density = = 0.5 in a simulated outbreak. B: Distribution of the posterior probability of direct transmission inferred by our algorithm for pairs of individuals in which a link existed in the simulation (red) and pairs of individuals which were not linked (blue).

Out of the *n* = 100 sampled individuals, only 53 were infected by another sampled individual; for the majority of these links, our algorithm inferred the existence of the link with high posterior probability, with only nine pairs being given a probability < 0.2 and 15 pairs being given a probability < 0.5 (Figure 2B, red curve). Conversely, for the 9847 pairs of sampled individuals for which a link did not exist in the simulated data, most were given a very small probability of a link in the posterior distribution of transmission trees, with only nine pairs being given a probability > 0.5 (Figure 2B, blue curve). If we consider 0.5 as the probability threshold for when transmission was inferred, our method had a specificity (true negative rate) of 99.9% and a sensitivity (true positive rate) of 72%. The area under the receiver operating characteristic (ROC) curve was 98.97%. These results demonstrate that in this specific example our algorithm was able to infer the correct transmission links with high accuracy, in spite of having information about only a proportion π = 0.5 of infected individuals. It should be noted that this application represents a best case scenario, since the phylogeny is known exactly, whereas for real epidemiological investigations it would need to be inferred from sequences, adding noise and uncertainty.

### Evaluation of performance using multiple simulated datasets

We repeated the simulation described above for values of the sampling density π varying from 0.1 to 1 by increments of 0.01, while leaving the reproduction number constant at *R* = 2. For each of the 90 simulated datasets, we applied our algorithm to estimate the values of both π and *R* (Figure 3). We found that the estimate of R remained fairly constant as it should, while the estimate of π increased as the correct value of π was increased. There was no sign of a bias in the estimates up to π = 0.6, but higher values of π were consistently underestimated, with the value of *R* being slightly overestimated in compensation. We attribute this bias to the difficulty in assessing with certainty whether all cases have been sampled in a transmission chain, since there always remains a possibility that an unsampled individual may have acted as intermediate. This small bias also reflects our choice of prior for π, which was uniform between 0 and 1, and the fact that only 100 genomes were used in each simulation.

**Figure 3.**
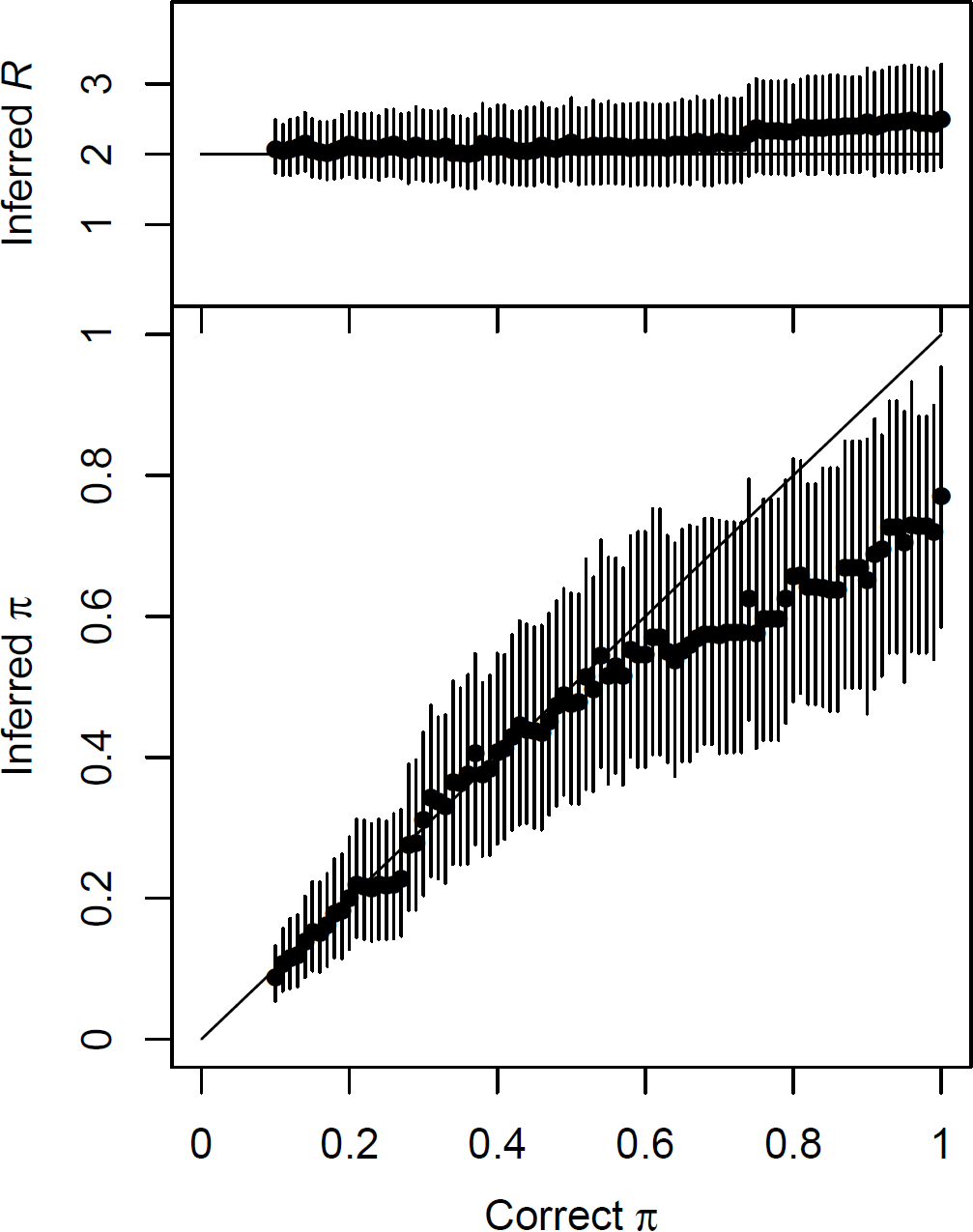
Inferred values of the reproduction number R (top) and the sampling proportion π (bottom) in simulated datasets for which the correct value of R is 2, and the correct value of π is increased from 0.1 to 1 (as shown on the x-axis). Dots represent the mean of the posterior sample and bars the 95% credibility intervals.

We also performed simulations in the converse situation where the sampling density was kept constant at π = 0.5 but the reproduction number was increased from *R* = 1 to *R* = 11 by increments of 0.1. For each of the 100 simulated datasets, our method was applied and the inferred values of π and *R* were recorded (Figure 4). Although there was once again a slight bias towards underestimating the sampling density π, its 95% credibility intervals always covered the correct value of π = 0.5. The inferred values of *R* were accordingly overall slightly upward biased, although they followed almost linearly the correct values used for simulation. The 95% credibility intervals for *R* almost always included the correct values. We conclude from these results that our algorithm performs well despite being tested in difficult situations, with only 100 sampled genomes, unknown proportions of unsampled cases, uninformative priors, and very large intervals of values being used in the simulations. A small outbreak with high sampling density and a larger outbreak with lower sampling density can often look similar, especially in the first stages of an ongoing outbreak, but our algorithm is able to distinguish between these two scenarios with good accuracy.

**Figure 4.**
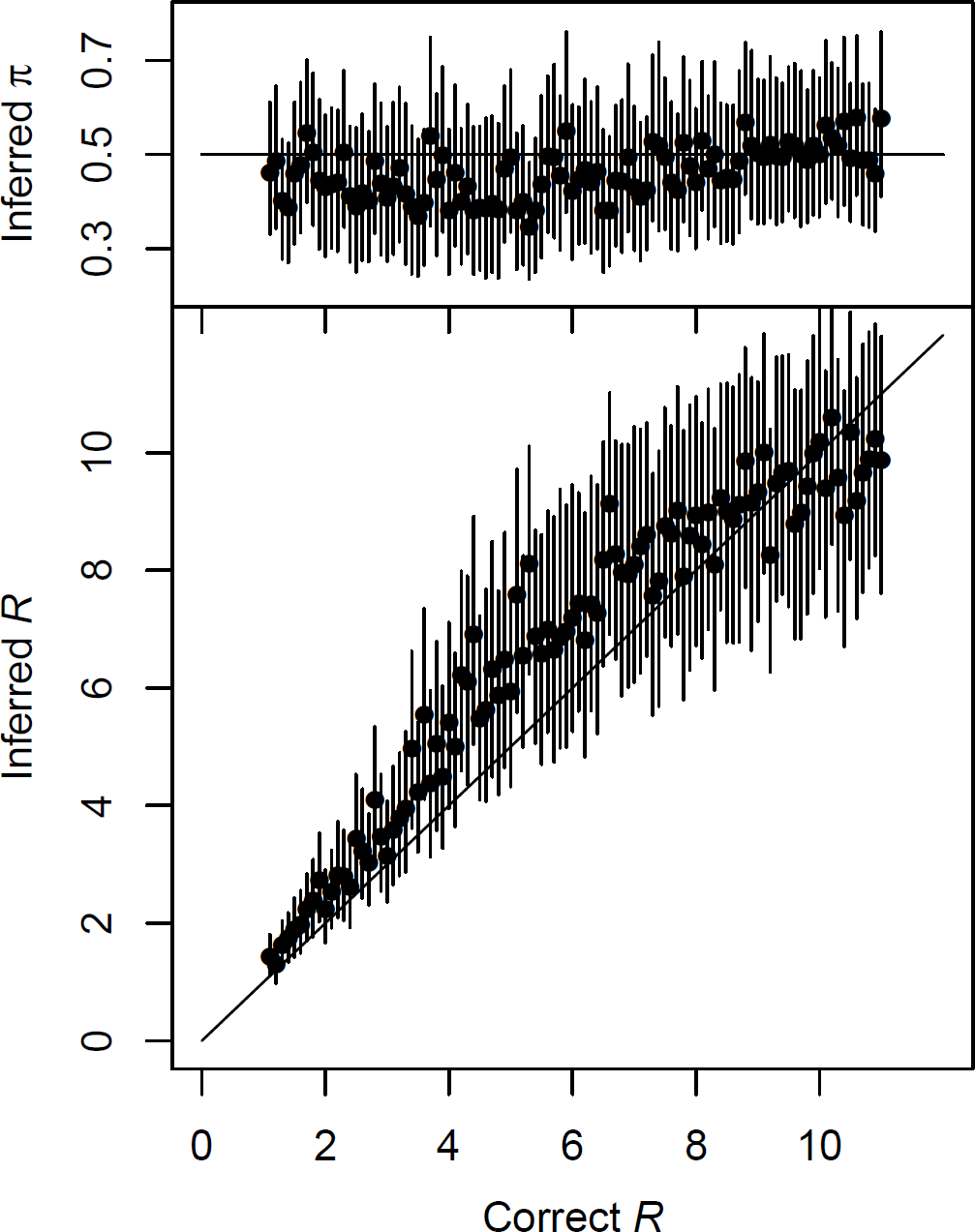
Inferred values of the sampling proportion π (top) and the reproduction number R (bottom) in simulated datasets for which the correct value of π is 0.5, and the correct value of R is increased from 1 to 11 (as shown on the x-axis). Dots represent the mean of the posterior sample and bars the 95% credibility intervals.

### Application to a *Mycobacterium tuberculosis* outbreak dataset

We applied the method to a previously reported tuberculosis outbreak (Roetzer et al., 2013). We used BEAST (Drummond et al., 2012) to infer a timed phylogeny from the published data (Figure S1). In determining the best priors for the densities of the times between becoming infected and infecting others (the generation time) and between becoming infected and becoming known to the health care system (sampling time), we considered both clinical aspects of tuberculosis disease and aspects of the epidemiological investigation. The outbreak lasted 13 years, during which active case finding was used to identify individuals with prior exposure to known cases. An early report on this outbreak (Diel et al., 2004) noted that many cases were identified for reasons not connected to their tuberculosis infection, such as presenting to health care with other symptoms, to obtain a health certificate, or to enter a detox program. We therefore used a Gamma distribution for the sampling time density, with a shape parameter 1.1 and rate 0.4. The generation time for tuberculosis should reflect a chance of relatively rapid progression from infection to active disease and hence to the opportunity to infect others, but also a possibility of infection leading to a long latent period before progression (Barry et al., 2009). We therefore used a Gamma function with shape parameter 1.3 and rate parameter 0.3 for the generation time density. We ran 100,000 MCMC iterations. The MCMC traces are shown in Figure S2.

Figure 5 shows the consensus transmission network for the real-world tuberculosis outbreak (Roetzer et al., 2013) and Figure 6 shows the inferred numbers of unsampled cases along with the reported cases through time. While most cases were sampled, reflecting a robust public health investigation, we estimate that early in the outbreak, several unsampled individuals were contributing to transmission. During this period, the two major clades of the phylogeny diverged. Figure 6A recapitulates the two major waves of the outbreak - an early peak around 1998 and a second pulse from 2005 onwards - each with a small portion of inferred unsampled cases. While the number of unsampled individuals was small, the method does allocate key transmission events to unsampled cases, particularly early in the outbreak, suggesting that screening and investigation earlier in the outbreak was not as comprehensive as it eventually became. This is to be expected, as outbreak management efforts typically intensify as the number of cases grows.

**Figure 5.**
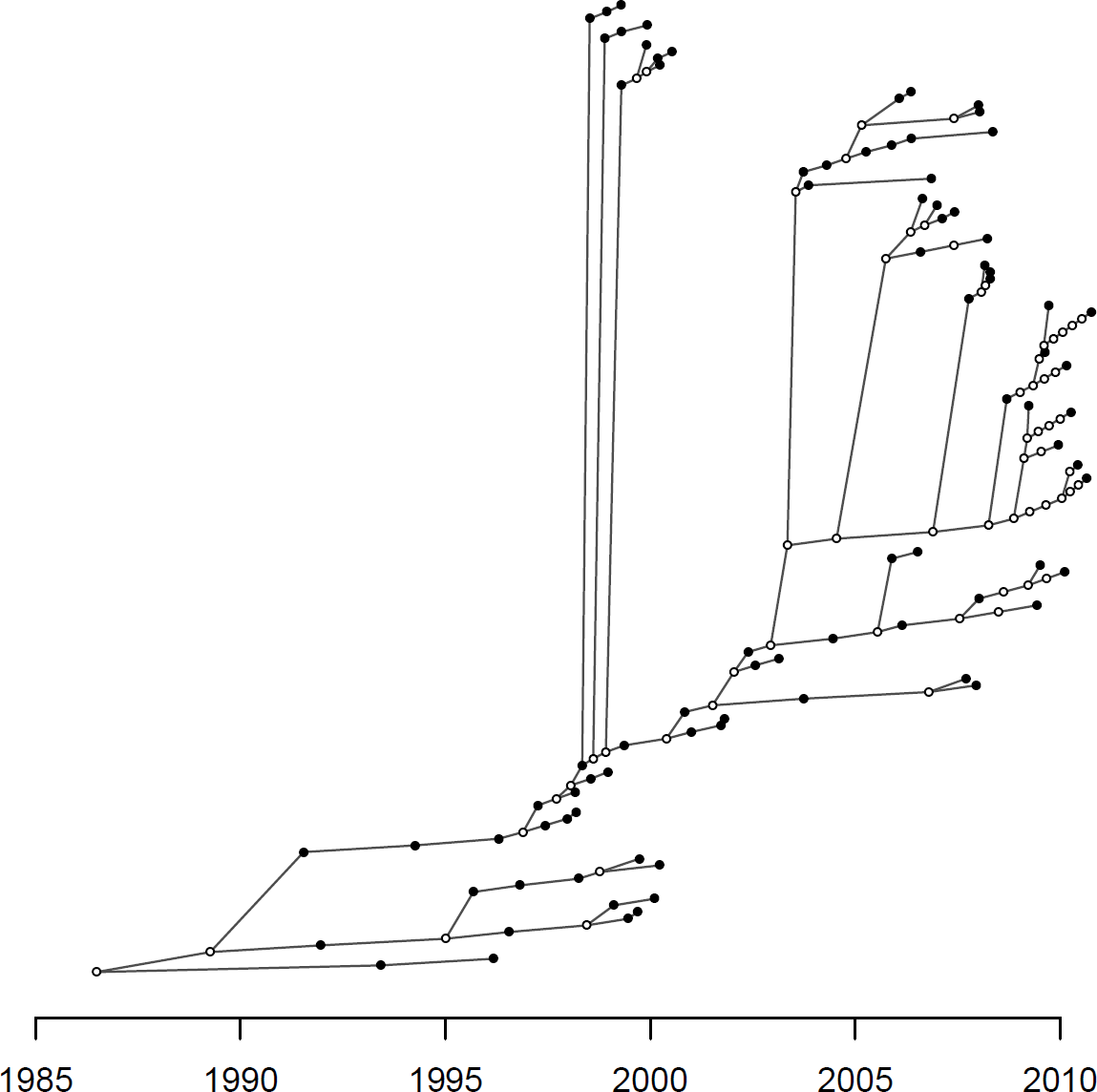
Consensus transmission tree for the tuberculosis outbreak. Filled dots represent sampled individuals and unfilled dots represent unsampled inferred individuals.

**Figure 6.**
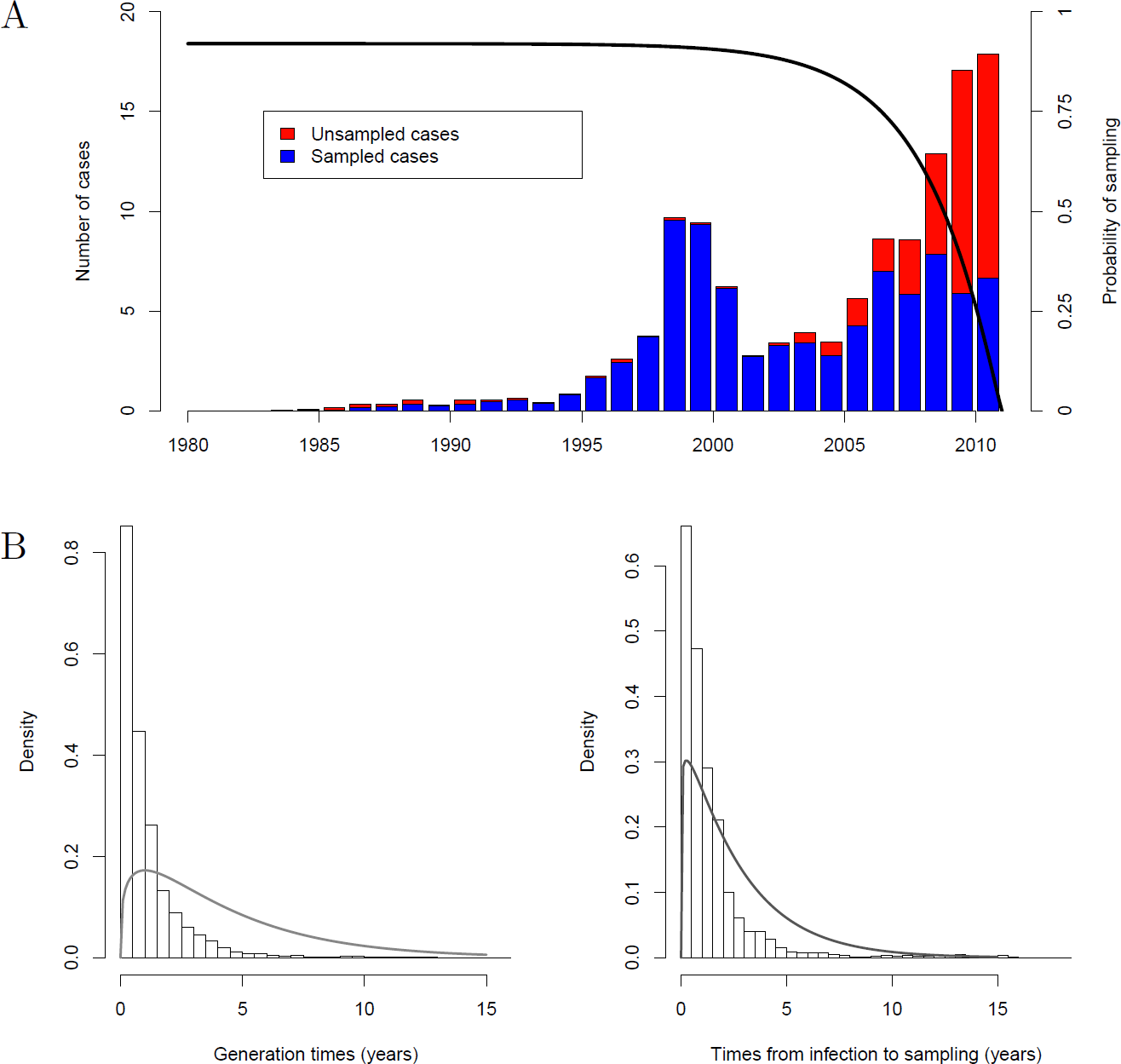
A: Outbreak plot showing the numbers of sampled and unsampled cases through time in the posterior sample of transmission trees. While the posterior estimate of π is 0.93, predicting that cases would eventually be detected with high probability, in the time period just before sampling ended, the inferred transmission trees contain a number of unsampled cases. The solid line represents the probability of sampling cases as a function of their infection time, given that observation stops at T = 2011. B: Posterior generation times and times between infection and sampling. Bars show histograms of the posterior quantities and solid lines show the related prior densities.

Figure 6B shows the posterior times between an individual becoming infected and infecting others - the generation time - and the posterior time intervals between infection and sampling - the infectious period, with priors shown in grey. Our observed generation times are variable, which reflects the clinical history of tuberculosis - an infection that can progress rapidly to active, infectious disease or that can have a asymptomatic, non-infectious latent period of variable length. We used a gamma function as a prior, with mode strictly greater than 0, but the posterior generation times have a mode of 0, suggesting a relatively high portion of those who go on to infect others have a rapid progression to from infection to active disease. It is important to note that the posterior generation times are only an indicator of the inferred natural history of tuberculosis *among those with active disease who were sampled*; individuals who were infected but did not progress to active disease and those who never presented to care and were not sampled do not appear in the dataset, and those who did not infect others do not appear in the cases behind the inferred generation times. The mean posterior generation time was 1.0 years with a standard deviation of 1.36 years. The posterior times between becoming infected and becoming known to health authorities also differ from the prior assumption; they have a mean of 1.4 years and standard deviation of 2 years. Sampling times are distinct from the prior but are affected by a change in the prior assumption.

Where inferred infectors are sampled cases with associated clinical and/or epidemiological data, an advantage of our approach is that it allows comparison of the relative contributions of different groups of individuals to the burden of transmission. Figure S3 shows the inferred per-case transmission stratified by several characteristics of the cases (Roetzer et al., 2013): individuals’ AFB smear status (a measure of how many bacilli are found in their sputum, if any), HIV status, abuse of alcohol or other drugs, and whether the individual had a permanent domestic residence. Our method did not detect significant differences in secondary infections arising from smear-positive and -negative cases, between substance users and non-substance users, and between stably or transiently housed individuals. However, consistent with the fact that HIV-positive patients tend to be less infectious with tuberculosis, we find that HIV-positive individuals transmitted somewhat fewer cases on average than HIV-negative individuals. Due to the small number of HIV-positive cases - only five individuals were HIV-positive in this data-the estimates are much more variable than the estimates for HIV-negative cases. Many more clinical or demographic factors might impact transmissions, such as the presence of cavitary disease and the reported number of social contacts, but these data were unavailable for the present analysis.

Results in Figure S3 do not reflect differences in transmission rates given contact with others, because we do not know about exposures that did not result in infection. We also do not have information about behaviours that might modulate transmission. For example, if smear-positive cases sought and obtained treatment more rapidly than smear-negative cases, or were more unwell and had more limited activities, their transmission rate per contact could be higher than their smear-negative counterparts but they might still contribute fewer onward transmissions. The posterior sampling density is π = 0.93 with a standard deviation of 0.05, consistent with a very densely-sampled outbreak in a high-resourced setting with good case finding. Posterior estimates of π depend somewhat on priors for σ.

## Discussion

We have described a new methodology for reconstructing who infected whom based on genomic data from an infectious disease outbreak. The novelty of this approach, which extends our earlier work in the area, is that it now accounts for both the possibility of some cases not having been sampled and the possibility that more cases may occur in the future. Addressing these issues overcomes key hurdles in using genomic data to reconstruct disease transmission events during a real-time public health response. In these situations, a case may not be sequenced due to a lack of clinical specimen or otherwise sequenceable material, while cases might go unsampled for various reasons, including subclinical, or asymptomatic, infections for which an individual may not seek care or a diagnosis in another jurisdiction. Furthermore, following early proof-of-concept retrospective studies, genomic epidemiology is now being used to prospectively understand outbreaks, as in the recent outbreak of Ebola (Gire et al., 2014). Allowing inference before the end of the outbreak turns our method into a real-time, actionable approach.

Our methodology is based on an explicit transmission model which makes a number of assumptions, some of which could be relaxed if required by specific applications. A first example is the fact that in our model the reproduction number *R* remains constant throughout the outbreak, whereas in many situations the reproduction number varies over time and quantifying these variations is of great epidemiological importance (Cori et al., 2013). This could be incorporated in our method relatively easily, for example assuming stepwise changes or some predetermined parametric function for *R(t)*. A second example concerns the observation of cases, which we assumed to happen with probability π(*t*) for an individual infected at time *t* with π(*t*) reflecting the impossibility of observing cases happening after the time *T* when observation stops and the lower probability of observing cases soon before *T* (Equation 8). It is often difficult in epidemiological studies to know the real function π(*t*), but in situation where for example surveillance did not start before a certain date, the function π(*t*) we used here could be updated to reflect this. There are also a few other assumptions in our model that would be more difficult to relax, such as the complete transmission bottleneck which considers that only a single pathogen variant is transmitted from the donor to the recipient of each transmission event.

A key feature of our methodology is that it proceeds in two steps - first, genomics data is used to reconstruct a phylogenetic tree, and second, likely transmission events given the phylogeny are inferred. There are both advantages and disadvantages to this approach, compared to the more theoretically accurate joint inference of phylogenetic and transmission trees (Hall et al., 2015). Our two-step approach makes it difficult to pass the uncertainty in the phylogenetic reconstruction on to the transmission analysis. This is especially relevant if the time-labelled phylogeny is inferred not using a point estimation procedure (Fourment and Holmes, 2014; To et al., 2016), but rather with a Bayesian sampling method (Drummond et al., 2012; Bouckaert et al., 2014). In this case, applying the transmission analysis separately to a sample of trees from the phylogenetic posterior can help account for uncertainty (Didelot et al., 2014). However, two problems remain: how to choose the tree prior in the phylogenetic reconstruction and how to combine the results from the separate transmission analyses. A solution may be to see the phylogenetic trees sampled in the first step as coming from a biased distribution, and correcting for this using importance sampling in the second step, such that the separate transmission analyses are correctly aggregated and the prior used in the first step is nullified (Meligkotsidou and Fearnhead, 2007). On a more positive note, it should be noted that our two-step approach has significant advantages both computationally and conceptually. Computationally, we were able to analyse outbreaks with hundreds of cases in a matter of hours. Conceptually, working with a fixed phylogeny allows us to explore much more complex models for transmission trees, such as the partially sampled and ongoing scenarios. To date, no other transmission inference approaches handle these difficult scenarios.

We have previously applied earlier versions of our approach to understanding a complex tuberculosis outbreak in a largely homeless Canadian population (Didelot et al., 2014; Hatherell et al., 2016), showing how reveals key individuals contributing to transmission and how its ability to time infection events can be used to declare a waning tuberculosis outbreak truly over. Here, we demonstrate our new methodology’s ability to identify unsampled cases. Finding such cases is critically important for tuberculosis control - not only does it allow us to seek out these individuals and connect them with treatment, but it allows us to extend our case-funding efforts to include a larger proportion of potentially exposed individuals. In our present analysis of the Hamburg dataset, we found that the generation time was relatively rapid, with the majority of infected individuals progressing to active disease and infecting others doing so within two years, with many progressing to active disease almost immediately. This is important data for outbreak management - if borne out by further reconstructions, it suggests a bound for the time over which an individual who has been exposed to tuberculosis should be followed up.

In conclusion, we present a new method for the automated inference of person-to-person disease transmission events from pathogen genomic data, one which accounts for the complex and variable nature of sampling cases during an outbreak. When coupled to the routine genomic surveillance of key pathogens now in place at many public health agencies, such as Public Health England’s new genomic approach to tuberculosis diagnosis and laboratory characterisation (Pankhurst et al., 2016), our method has the potential to rapidly suggest the contact network underlying an outbreak. Given the significant resources associated with a contact investigation, any tool that can quickly assist in prioritising individuals for followup is an important contribution to the public health domain.

## Acknowledgments

This work was supported by the UK National Institute for Health Research Health Protection Research Unit in Modelling Methodology at Imperial College London in partnership with Public Health England (grant HPRU-2012-10080 to XD) and the UK Medical Research Council (grant MR/N010760/1 to XD). JG holds a Canada Research Chair in Public Health Genomics and a Michael Smith Foundation for Health Research Scholar Award. The funders had no role in study design, data collection and interpretation, or the decision to submit the work for publication.

